# Cold sensitivity of the SARS-CoV-2 spike ectodomain

**DOI:** 10.1101/2020.07.12.199588

**Authors:** Robert J Edwards, Katayoun Mansouri, Victoria Stalls, Kartik Manne, Brian Watts, Rob Parks, Katarzyna Janowska, Sophie M. C. Gobeil, Megan Kopp, Dapeng Li, Xiaozhi Lu, Zekun Mu, Margaret Deyton, Thomas H Oguin, Jordan Sprenz, Wilton Williams, Kevin Saunders, David Montefiori, Gregory D. Sempowski, Rory Henderson, Munir Alam, Barton F. Haynes, Priyamvada Acharya

**Author notes:** These authors contributed equally (Robert J Edwards, Katayoun Mansouri, Victoria Stalls, Kartik Manne). To whom correspondence should be addressed, Correspondence to: Robert J Edwards and Priyamvada Acharya.

## Abstract

The SARS-CoV-2 spike (S) protein, a primary target for COVID-19 vaccine development, presents its Receptor Binding Domain in two conformations: receptor-accessible “up” or receptor-inaccessible “down” conformations. Here, we report that the commonly used stabilized S ectodomain construct “2P” is sensitive to cold temperature, and that this cold sensitivity is resolved in a “down” state stabilized spike. Our results will impact structural, functional and vaccine studies that use the SARS-CoV-2 S ectodomain.

---

The spike (S) protein of SARS-CoV-2 mediates receptor binding and cell entry and is a key target for vaccine development efforts. Stabilized S ectodomain constructs have been developed that mimic the native spike, bind ACE-2 receptor^1,2^, and present epitopes for neutralizing antibodies on their surface^3–6^. The “2P” S-ectodomain construct examined here incorporates residues 1-1208 of the SARS-CoV-2 spike, with two proline mutations in C-terminal S2 domain designed to stabilize the pre-fusion spike conformation, a C-terminal foldon trimerization motif, and a mutation that abrogates the furin cleavage site^1^ (Figure 1a). This and similar constructs are being widely used for structural biology and vaccine studies^1–3,7,8^. Purified S ectodomain proteins^9^ are assessed for quality control by SDS-PAGE, size exclusion chromatography (SEC), differential scanning fluorimetry (DSF)^10^, and negative stain electron microscopy (NSEM). NSEM has proved especially important because it reveals the threedimensional structural integrity of individual molecules, allowing us to discriminate between preparations that looked similar by bulk methods such as SDS-PAGE and SEC (Extended Data Figure 1). The observed variability in the spike fraction between preparations suggests a fragile S ectodomain, and measures to overcome the issue have been previously reported^11,12^. Here we link the apparent fragility of spike ectodomain 2P to its rapid denaturation upon storage at 4 °C (Figure 1 b-e).

**Figure 1.**
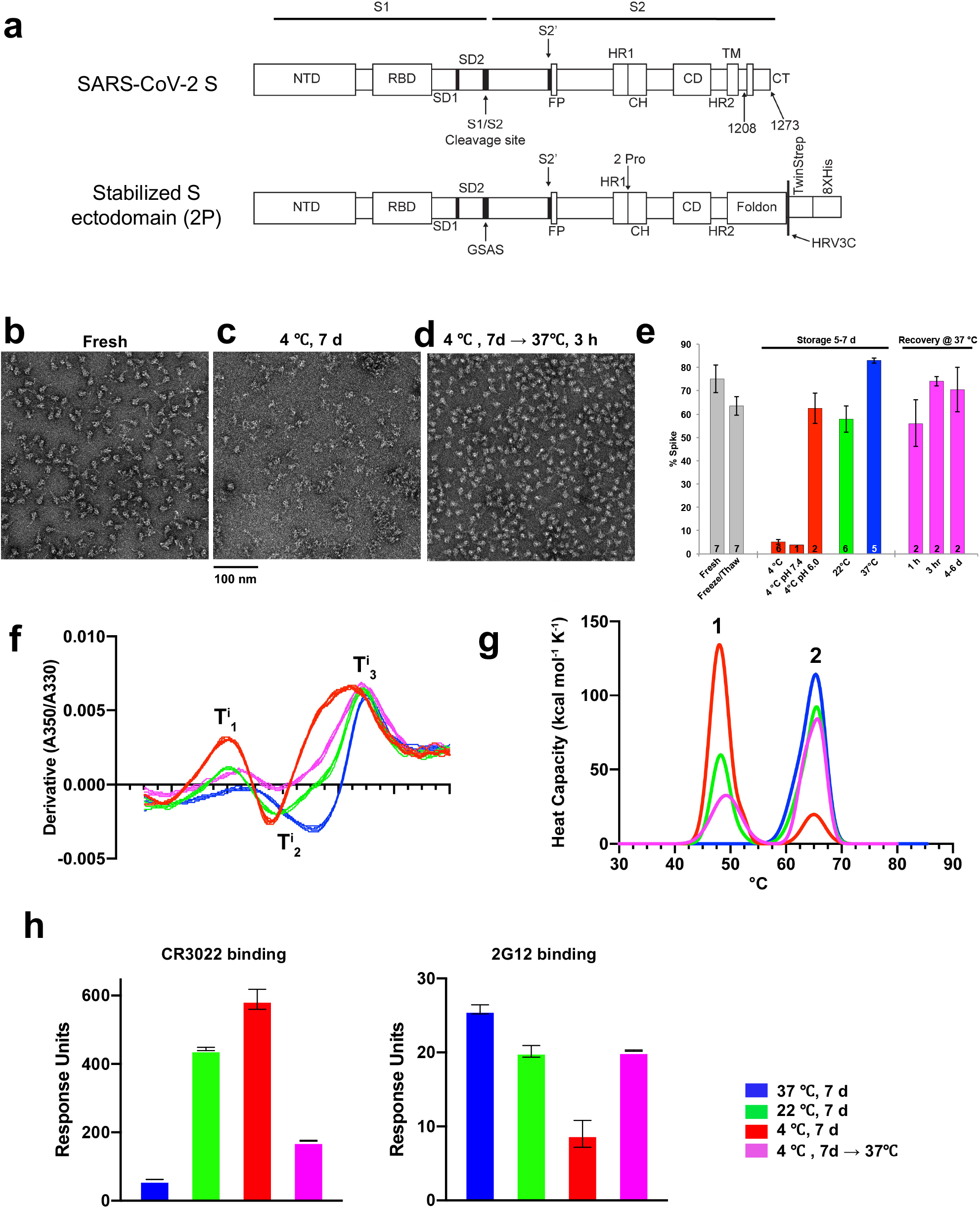
Temperature-dependence of the SARS-CoV-2 S ectodomain. **(a)** Schematic of the SARS-CoV-2 spike (top) and a furin cleavage-deficient, soluble, stabilized ectodomain construct (bottom). **(b-d)** Representative NSEM micrograph from **(b)** a freshly prepared sample of the S ectodomain, **(c)** the same sample after storing the protein for one week at 4 °C, **(d)** the same sample after storing the protein for one week at 4 °C followed by a 3-hour incubation at 37 °C. **(e)** Bar graph summarizing NSEM results on S ectodomain samples stored under different conditions. From left to right, bars show spike percentage in a fresh sample of spike; after the spike undergoes a single freeze-thaw cycle; after it has been incubated for 5-7 days at 4 °C (red), 22 °C (green) or 37 °C (blue); and spike samples that were stored for 1 week at 4 °C, then incubated at 37 °C for 1 hour, 3 hours, or 4-6 days. Solid bars indicate averages, with number of samples indicated at the bottom of each column. The error bars indicate standard error of the mean, except when N=2, where the error bars indicate range. **(f)** DSF profiles obtained by following changes in intrinsic fluorescence upon applying a thermal ramp to the sample and expressed as the first derivative of the ratio between fluorescence at 350 nm and 330 nm. Maxima and minima indicate inflection temperatures, T. For each data point, 5 overlaid curves are shown indicating technical replicates. **(g)** DSC profiles, shown as representative of two technical repeats. **(h)** Antibody CR3022 IgG (left) and 2G12 IgG (right) binding to spike stored at different temperatures measured by SPR. Data for spike samples measured after a 1-week incubation at 37, 22, and 4 °C, are shown in blue, green, and red respectively; sample stored 1 week at 4 °C and then incubated for 6 hours at 37 °C shown in magenta. During the SPR run the sample chamber was maintained at temperatures of 37 °C, 22 °C and 8 °C, for the 37 °C, 22 °C and 4 °C incubated samples, respectively. The binding experiments were carried out at 25 °C. Error bars are derived from 3 technical repeats. The data shown are representative of at least 5 independent experiments. 2 independent repeats using separate protein lots are shown in Supplementary Figure 2.

We followed the structural, biophysical and antigenic features of the S ectodomain stored under different temperatures conditions (Figure 1, Extended Data Figures 2-6). The 2P spike was produced in 293F cells at 37 °C and the purification was performed at room temperature and completed within 6-8 hours (Supplementary Figure 1a-c). Representative NSEM micrographs obtained of the 2P spike incubated at different temperatures and shown (Figure 1b-d, Supplementary Figure 1d-e), and results summarized in Figure 1e. Freshly prepared spike samples assessed on the same day they were purified showed 75% well-formed spikes on average (Fig 1b,e); this fraction slightly decreased to 64% after one cycle of freeze/thaw and to 59% by storage at room temperature (22 °C) for 5–7 days (Supplementary Figure 1d,e); and it was substantially reduced to 5% after storage at 4 °C for 5–7 days (Fig 1c,e). In contrast, after 1-week storage at 37 °C, we observed 83% well-formed spike. Furthermore, well-formed spike could be recovered after the spike was stored at 4 °C for a week with a 3-hour incubation at 37 °C to ~75% (Figure 1d,e); no further improvement was observed after longer incubations at 37 °C (Figure 1e) which actually led to slight aggregation (Supplemental Figure 1e). SDS-PAGE analysis (Supplementary Figure 1) of 2P spike samples stored at different temperatures does not indicate any appreciable degradation of the spike, although we do observe evidence of higher molecular weight species in spike stored at 37 °C for a week (Supplemental Figure 1). Similar to our NSEM results, we also see substantial differences in the quality and dispersion of protein in cryo-EM grids depending on the temperature the specimen was stored (Extended Data Figure 6). In summary, these data show that the 2P spike denatured upon storage for at 4 °C for a week, and this denaturation could be at least partially reversed by incubating the sample at 37 °C following cold-storage.

The S ectodomain was reported to exhibit conformational changes in response to pH^12^, so we asked whether the spike denaturation observed was due to temperature-dependent pH changes at 4 °C in the T ris pH 8.0 buffer we used^13^. A Tris buffer solution that measures pH 8.0 at 25 °C would measure ~pH 8.6 at 4 °C. We performed SEC purification of a 2P spike preparation into MOPS buffer pH 7.4. MOPS has a smaller temperature dependence compared to Tris^13^, and is expected to change only slightly to pH 7.42 at 4 °C. Cold-storage at pH 7.42 (MOPS buffer) reduced the spike fraction to 4%, similar to the average of 5% for cold-storage in Tris buffer (Figure 1e). Thus, the primary cause of spike denaturation appears to be the temperature change and not the pH shift within this range. We also tested the temperature effect at acidic pH and found that incubating the spike at 4 °C in MES, pH 6 buffer reduced but did not eliminate the cold-sensitivity (Fig 1e).

We next examined whether the observed cold sensitivity is a property of the bulk protein in solution, rather than an artifact of the NSEM sample preparation. First, we used a rapid DSF assay^10^ that measures changes in the intrinsic fluorescence of the protein as a thermal ramp is applied. The changes in fluorescence signal indicate transitions in the folding state of a protein, the temperature at which a transition occurs is called the inflection temperature (Ti). Distinct profile shifts were observed for spike samples stored at different temperatures, with Ti shifted toward lower temperatures for spike samples stored at 4 °C, indicating lower protein stability, compared to when the spike was stored at 22 °C or 37 °C (Figures 1f). Next, we measured melting temperatures (Tm) using differential scanning calorimetry (DSC) (Figure 1g and Extended Data Figure 3): after one-week storage at 37 °C, the spike sample featured an asymmetric unfolding transition with T_m_ of 65.5 °C. After one-week storage at 22 °C, we observed a second low-Tm transition at 48.2 °C. After one-week storage at 4°C, we observed a similar two-peak profile with a markedly more pronounced low-T_m_ transition (T_m_= 48.4 °C). Upon returning the 4 °C sample to 37 °C for 3 hours prior to analysis, we observed an amplitude reduction of the low-T_m_ transition (T_m_= 49.2 °C) and a corresponding amplitude increase in the high-T_m_ transition (T_m_= 66.0 °C). Thus, the DSC results confirm that storage at 4 °C destabilizes the spike compared to samples stored at 22 °C or 37 °C, and that returning the destabilized spike to 37 °C for 3 hours substantially restores its stability, although the presence of the low-Tm peak suggests that the recovery is partial.

We next tested the effects of cold-induced spike instability on antibody and receptor binding by SPR and ELISA (Figures 1h and Extended Data Figures 4-5, Supplementary Figure 2). We found that spike stored at 4 °C showed higher binding to antibody CR3022 (Figure 1h, Extended Data Figure 4, Supplementary Figure 2) and the receptor ACE-2 (Extended Data Figure 4), which both require an “up” RBD conformation for stable binding^1,9,12^. In contrast, cold-storage reduced binding to antibody 2G12 (Figure 1h, Extended Data Figure 4, Supplementary Figure 2), which recognizes a quaternary glycan epitope in the S2 subunit^14^ indicating loss of quaternary structure during cold storage. We also tested two antibodies isolated from a COVID-19 convalescent donor (Supplementary Figures 4-5), with epitopes mapped to the ACE-2 binding site (AB712199) and to the S2 region of the spike (AB511584) (Supplementary Figure 4). Both antibodies showed different binding profiles depending on the temperature at which the spike was stored, highlighting the importance of accounting for this cold-sensitive behavior of the S ectodomain for obtaining consistent results in serology assays when using the 2P spike or similar constructs. We next tested CR3022 and 2G12 binding to freshly prepared 2P spike sample and compared it to a sample that was flash frozen in liquid N2, stored in single-use aliquots at −80 °C and thawed by incubating at 37 °C Supplementary Figure 3). We found that freeze-thaw increased CR3022 binding to the 2P spike relative to the fresh sample, although incubation at 37 °C of the thawed sample for 20 min reduced binding close to the levels observed in the fresh sample. Based on these data, in order to obtain consistent results in binding studies, we recommend flash-freezing spike samples in single use aliquots, thawing at 37 °C followed by a brief (~20 min) incubation at 37 °C before using the 2P spike in binding studies.

These results suggested that cold-induced denaturation of spike was associated with increased RBD-exposure of the S ectodomain, and we thus asked whether a “down” state stabilized S ectodomain might be resistant to cold-induced denaturation (Figure 2). We included in this analysis a variant that combined the recently described rS2d^9^ and HexaPro mutations^11^ (Figure 2a,b). This new variant, named rS2d-HexaPro, showed the highest production yields of the all-RBD-down spike constructs (Figure 2c). Similar to the rS2d construct^9^, rS2d-Hexapro was shown by NSEM to be 100 % in a 3-RBD-down conformation (Figure 2d,e, Extended Data Figure 7 and Supplementary Movie 1). Both HexaPro and rS2d-HexaPro appeared more resistant to cold-induced denaturation (Figure 2f) than the 2P spike (Figure 1e), with rS2d-HexaPro showing a higher intact spike percentage than HexaPro (Figure 2f). The rS2d mutations (D985C, S383C) introduce an inter-protomer disulfide bond between the RBD and the S2 domain, which we suggest stabilizes the spike and prevents its cold-induced denaturation. An independent study also reported that the 985C-383C disulfide bond, as well as an introduced disulfide between residues 987 and 413, prevented denaturation of the spike when stored at 4 °C^15^. These data suggest that inter-promoter disulfide linkages may be a general strategy for stabilization of the 2P spike ectodomain and preventing its cold-induced denaturation. We also included two glycan-deleted mutants, 2P-N165A and 2P-N234A^16^, that favor the RBD “up” or “down” conformation, respectively (Figure 2f), and both mutants showed substantial reduction of spike percentage after incubation at 4 °C (Figure 2f), Thermostability and binding studies (Figures 2g,h) further confirmed that both HexaPro and rS2d-HexaPro were more resistant to cold destabilization compared to the 2P version (Figure 1). ACE-2 binding to rS2d-HexaPro was substantially reduced for samples stored at all temperatures, as expected for a spike construct fixed in an RBD-down conformation. Similarly, binding to 2G12 remained same for the rS2d-HexaPro construct irrespective of the storage temperature of the 2P spike (Figure 1h), showing that the rS2d-Hexapro structure remains unperturbed upon 4 °C storage. For HexaPro, binding to ACE-2 was higher and 2G12 binding was reduced with 4°C-stored protein compared to 37 °C-stored samples, showing that the HexaPro structure remained susceptible to perturbation at lower temperatures.

**Figure 2.**
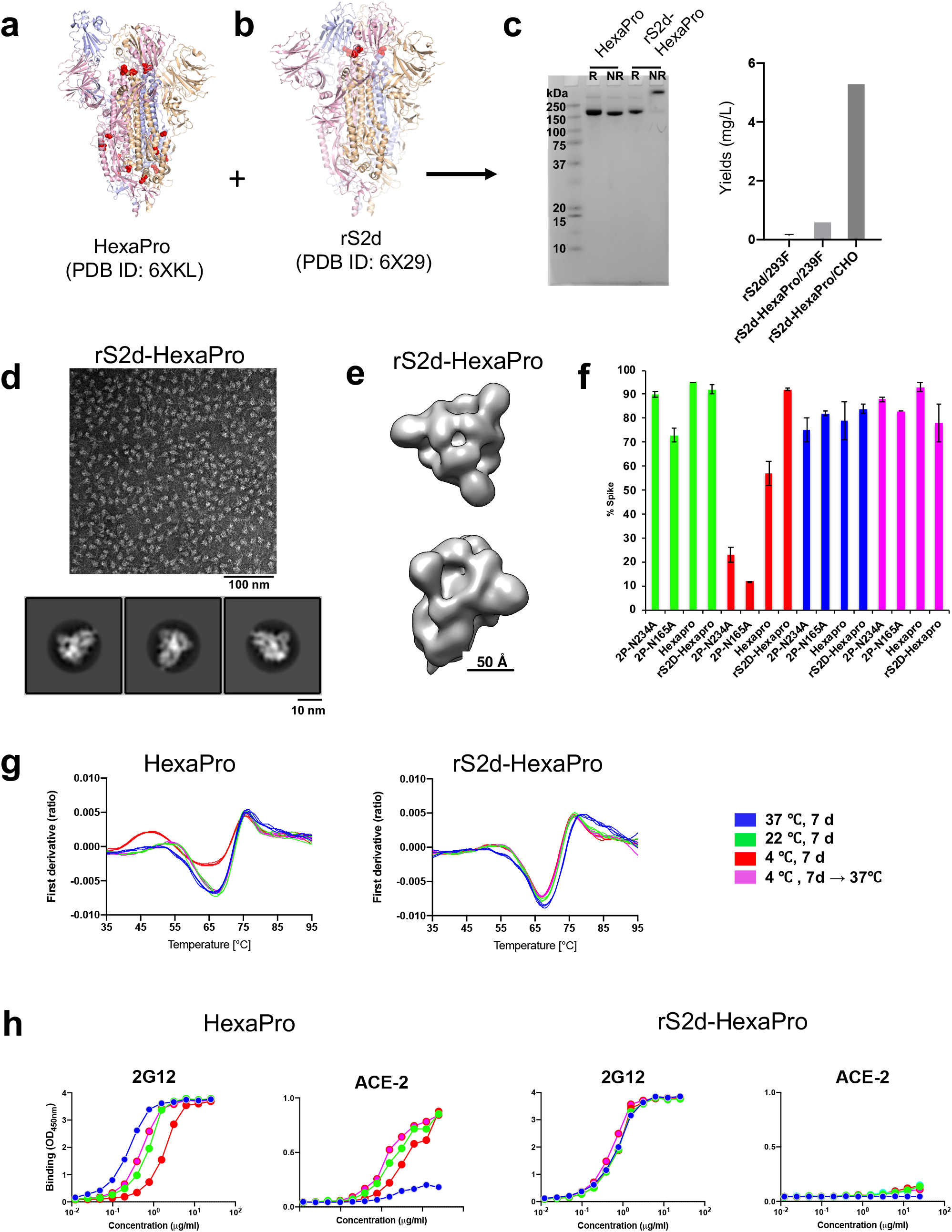
Engineered SARS-CoV-2 spike variant, rS2d-HexaPro, is resistant to temperature-dependent structural changes. **(a-b)** Structures of **(a)** HexaPro showing a 1-RBD-up conformation (PDB ID: 6X29) and **(b)** rS2d (PDB ID: 6XKL) showing a all-RBD-down conformation. **(c)** *(left)* SDS-PAGE. Lane 1: Molecular weight marker, Lanes 2 and 3: HexaPro, Lane 4 and 5: rS2d-HexaPro. R= Reducing, and NR = Non-reducing conditions. *(right)* Bar graph summarizing protein yields in mg/L of culture for, from left to right, rS2d produced in 293F cells, rS2d-HexaPro produced in Freestyle 293F cells, and rS2d-HexaPro expressed in CHO cells. **(d)** *(top)* Representative NSEM micrograph from a preparation of rS2d-Hexapro and *(bottom)* 2D class averages. **(e)** 3D reconstruction of rS2d-Hexapro obtained from NSEM data. **(f)** Bar graph summarizing results from NSEM on spike variants stored at different temperatures. Error bars shown are derived from at least two independent experiments with dfferent protein lots, and represent standard error of mean where N ≥ 3 and range where N=2. **(g)** DSF profiles for Hexapro (left) and rS2d-Hexapro (right). **(h)** Binding of 2G12 and ACE-2 to Hexapro (left) and rS2d-Hexapro (right) measured by ELISA. Serially diluted spike protein was bound in individual wells of 384-well plates, which were previously coated with streptavidin. Proteins were incubated and washed, then 2G12 at 10 μg/ml or ACE-2 with a mouse Fc tag at 2 μg/ml were added. Antibodies were incubated, washed and binding detected with goat anti-human-HRP.

Overall, our results demonstrate the cold sensitivity of the furin-cleavage deficient SARS-CoV-2 ectodomain, and its impact on antibody and receptor binding. We have also shown that this cold destabilization is substantially reduced in the HexaPro spike and further reduced by stabilization of the S ectodomain in a disulfide-locked “down” position.

## Acknowledgements

This work was supported by NIH NIAID extramural project grants R01 AI145687 (P.A.), and AI058607 (G.S.), funding from the Department of Defense HR0011-17-2-0069 (G.S.) and a contract from the State of North Carolina Pandemic Recovery Office through funds from the Coronavirus Aid, Relief, and Economic Security (CARES) Act (B.F.H.). This work utilized the DSC platform supported by the Duke Consortia for HIV/AIDS Vaccine Development (CHAVD) and the Titan Krios microscope in the Shared Materials and Instrumentation Facility at Duke University. The following reagent was deposited by the Centers for Disease Control and Prevention and obtained through BEI Resources, NIAID, NIH: SARS-Related Coronavirus 2, Isolate USA-WA1/2020, NR-52281. MN and PRNT assays were performed in the Virology Unit of the Duke Regional Biocontainment Laboratory, which received partial support for construction from the National Institutes of Health, National Institute of Allergy and Infectious Diseases (UC6-AI058607).

## Author contributions

R.J.E., K.Mansouri., V.S. and P.A. discovered the effect of storage temperature on spike stability. R.J.E. led NSEM studies and established quantitative metrics for spike QC. K.Mansouri collected NSEM data and performed analyses. V.S. purified proteins and performed thermostability measurements. K.Manne purified proteins, performed SPR assays, prepared samples for NSEM, Tycho, ELISA, SPR and DSC measurements, and coordinated the study between the different research teams. B.W. performed DSC measurements. R.P., M.D. J.S. and W.W. performed ELISA assays. D.L., X.L., K.S. and G.S. isolated antibodies from convalescent patient. S.G. and K. J. purified proteins and performed thermostability assays. M.F.K purified proteins. Z.M. performed binding measurements. R.H. initiated the DSC experiments and provided the rS2d, 2P-N234A and 2P-N165A constructs prior to publication. M.A. supervised the DSC experiments. B.F.H. supervised ELISA experiments and antibody isolation. P.A. oversaw and led the study, and co-wrote the paper with R.J.E. and V.S. All authors reviewed and commented on the manuscript.

## Methods

### Protein expression and purification

SARS-CoV-2 ectodomain constructs were produced and purified^1,11^ as follows. A gene encoding residues 1-1208 of the SARS-CoV-2 S (GenBank: MN908947) with proline substitutions at residues 986 and 987, a “GSAS” substitution at the furin cleavage site (residues 682-685), a C-terminal T4 fibritin trimerization motif, an HRV3C protease cleavage site, a TwinStrepTag and an 8XHisTag was synthesized and cloned into the mammalian expression vector pαH. Plasmids were transiently transfected into either FreeStyle-293F cells using Turbo293 (SpeedBiosystems). Cells were grown in a 37 °C incubator with 9 % CO2 and 120 rpm rotary motion. Protein was purified on the sixth day post-transfection from filtered supernatant using StrepTactin resin (IBA), followed by SEC purification using a Superose 6 10/300 Increase column in nCoV buffer (2mM Tris, pH 8.0, 200 mM NaCl, 0.02% sodium azide). rS2d-Hexapro was also grown in CHO cells. Plasmids were transiently transfected into CHO cells using ExpiFectamine CHO Transfection Kit (ThermoFisher), and protein was purified one the sixth day post-transfection. All protein purification steps including Strep tag purification and SEC were performed at room temperature. The spikes were purified the same day that the cells are harvested and the purification is completed 6-8 hours. The purified protein is flash frozen and stored in −80 °C in single-use aliquots. Each aliquot is thawed and briefly incubated (~20 min) at 37 °C before use.

Antibodies were produced in Expi293 cells and purified by Protein A affinity. For ACE-2 constructs, the ACE-2 C-terminus was fused with either the human or mouse Fc region including C-terminal 6X His-tag on the Fc domain. ACE-2 with human Fc tag was purified by Protein A affinity chromatography, and ACE-2 with mouse Fc tag was purified by Ni-NTA chromatography.

### Negative-stain electron microscopy

Spike samples were pre-incubated (stored) for specified times in nCoV buffer at 4, 22 or 37 °C, then moved to room temperature for preparation of NSEM grids, which was complete in less than 5 min. Samples were diluted to 100 μg/ml with room-temperature buffer containing 20 mM HEPES pH 7.4, 150 mM NaCl, 5% glycerol and 7.5 mM glutaraldehyde, and incubated 5 min; then glutaraldehyde was quenched for 5 min by addition of 1M T ris stock to a final concentration of 75 mM. A 5-μl drop of sample was applied to a glow-discharged, carbon-coated grid for 10-15 s, blotted, stained with 2% uranyl formate, blotted and air-dried. Images were obtained with a Philips EM420 electron microscope at 120 kV, 82,000× magnification, and a 4.02 Å pixel size. The RELION program^17^ was used for particle picking, 2D and 3D class averaging.

### Thermostability assays

Thermostability of the S ectodomain samples were measured using DSF and DSC. Samples were purified and buffer exchanged into HBS buffer (10 mM HEPES, 150 mM NaCl, pH 7.4) by SEC on a Superose 6 10/300 column. DSF assay was performed using Tycho NT. 6 (NanoTemper Technologies). Spike variants were diluted (0.15 mg/ml) in HBS and run in triplicate. Intrinsic fluorescence was recorded at 330 nm and 350 nm while heating the sample from 35-95 °C at a rate of 30 °C/min. The ratio of fluorescence (350/330 nm) and inflection temperatures (Ti) were calculated by Tycho NT. 6.

DSC measurements were performed using the NanoDSC platform (TA instruments; New Castle, DE). Samples that had been incubated at 22 °C or 37 °C were purified by SEC at room temperature and the sample incubated at 4 °C was purified by SEC at 4 °C (Extended Data Figure 2), diluted to 0.2–0.3 mg/mL in HBS, and degassed for 15 min at room temperature prior to analysis. DSC measurements were performed immediately after SEC purification of the samples. DSC cells were conditioned with filtered, degassed HBS prior to sample loading. Protein samples were heated from 10 °C to 100 °C at 1 °C/min under 3 atm pressure using HBS as the reference buffer. The observed denaturation profiles were buffer subtracted, converted to molar heat capacity, baseline-corrected with a 6th-order polynomial, and fit with 2-4 Gaussian transition models, as needed, using the NanoAnalyze software (TA Instruments). The peak transition temperature (Tm) is reported as the temperature at the maximum observed heat capacity of each transition peak.

### Isolation of antibodies from COVID-19 convalescent donors

Human SARS-CoV-2 Spike antibodies Ab712199 and Ab511584 were isolated from a COVID-19 convalescent individual. Peripheral blood was collected following informed consent on a Duke University Medical Center approved Institutional Review Board protocol. Briefly, PBMC samples collected after the onset of the symptoms were stained and the memory B cells were sorted with SARS-CoV-2 Spike-2p probes. Antibody IgH and IgK/L genes were recovered from the single-cell sorted cells, cloned into human IgG1 constant region backbone, and purified by Protein A beads as previously described^18^.

### ELISA assays

Spike samples were pre-incubated at different temperatures then tested for antibody- or ACE-2-binding in ELISA assays as previously described^19,20^. Assays were run in two formats. In the first format antibodies or ACE-2 protein were coated on 384-well plates at 2 μg/ml overnight at 4 °C, washed, blocked and followed by two-fold serially diluted spike protein starting at 25 μg/ml. Binding was detected with polyclonal anti-SARS-CoV-2 spike rabbit serum (developed in our lab), followed by goat anti-rabbit-HRP (Abcam #ab97080) and TMB substrate (Sera Care Life Sciences #5120-0083). Absorbance was read at 450 nm. In the second format, serially diluted spike protein was bound in individual wells of 384-well plates, which were previously coated with streptavidin (Thermo Fisher Scientific #S-888) at 2 μg/ml and blocked. Proteins were incubated at room temperature for 1 hour, washed, then human mAbs were added at 10 μg/ml. Antibodies were incubated at room temperature for 1 hour, washed and binding detected with goat anti-human-HRP (Jackson ImmunoResearch Laboratories, #109-035-098) and TMB substrate.

Commercially obtained constructs of SARS-CoV-2 spike ectodomain (S1+S2 ECD, S2 ECD and RBD) (Sino Biological Inc cat# 40589-V08B1 and 40590-V08B respectively and RBD from Genescript cat# Z03483) were coated directly on 384-well plates at 2 μg/ml and incubated overnight at 4 °C. Plates were washed, blocked and human mAbs three-fold serially diluted from 100 μg/ml were added for 1 hour at room temp followed by washing. Binding was detected with Goat anti-human IgG-HRP followed by TMB substrate.

### Microneutralization assay

Live SARS-CoV-2 Microneutralization (MN) assays were adapted from Berry et al.^21^ In short, sera or purified antibodies are diluted two-fold and incubated with 100 TCID50 virus for 1 hour. These dilutions are used as the input material for a TCID50. Each batch of MN includes known neutralizing and non-neutralizing controls. Data are reported as the last concentration at which a test sample protects Vero E6 cells. Recently, a fluorescent infectious clone of SARS-CoV-2 was developed^22^.

### Plaque Reduction Neutralization Test (PRNT)

Live SARS-CoV-2 Plaque Reduction Neutralization Test (PRNT) was performed in an assay adapted from Berry et al.^21^ Briefly, two-fold dilutions of a test sample (serum, plasma, purified Ab) are incubated with 50 PFU SARS-CoV-2 for 1 hour (NR-52281; BEI Resources). The antibody/virus mixture was then used to inoculate Vero E6 cells (ATCC). Cultures were then incubated at 37°C, 5% CO_2_ for one hour. At the end of the incubation, 1 mL of a viscous overlay (1:1 2X DMEM and 1.2% methylcellulose) was added to each well. Plates were incubated for 4 days. After fixation, staining and washing, plates were dried and plaques counted. Known neutralizing and non-neutralizing antibodies/sera are included in this assay as positive and negative controls, respectively. Plaques from each dilution of each sample are counted and data are reported as the concentration at which 50% of input virus is neutralized.

### Pseudovirus Neutralization Assay

SARS-CoV-2 neutralization was assessed with Spike-pseudotyped viruses in 293T/ACE2 cells as a function of reductions in luciferase (Luc) reporter activity. 293T/ACE2 cells were kindly provided by Drs. Mike Farzan and Huihui Mu at Scripps. Cells were maintained in DMEM containing 10% FBS and 3 μg/ml puromycin. An expression plasmid encoding codon-optimized full-length Spike of the Wuhan-1 strain (VRC7480), was provided by Drs. Barney Graham and Kizzmekia Corbett at the Vaccine Research Center, National Institutes of Health (USA). The D614G amino acid change was introduced into VRC7480 by site-directed mutagenesis using the QuikChange Lightning Site-Directed Mutagenesis Kit from Agilent Technologies (Catalog # 210518). The mutation was confirmed by full-length Spike gene sequencing. Pseudovirions were produced in HEK 293T/17 cells (ATCC cat. no. CRL-11268) by transfection using Fugene 6 (Promega Cat#E2692) and a combination of Spike plasmid, lentiviral backbone plasmid (pCMV ΔR8.2) and firefly Luc reporter gene plasmid (pHR’ CMV Luc)^23^ in a 1:17:17 ratio. Transfections were allowed to proceed for 16-20 hours at 37°C. Medium was removed, monolayers rinsed with growth medium, and 15 ml of fresh growth medium added. Pseudovirus-containing culture medium was collected after an additional 2 days of incubation and was clarified of cells by low-speed centrifugation and 0.45 μm micron filtration and stored in aliquots at −80°C. TCID_50_ assays were performed on thawed aliquots to determine the infectious dose for neutralization assays (RLU 500-1000x background, background usually averages 50-100 RLU).

For neutralization, a pre-titrated dose of virus was incubated with 8 serial 3-fold or 5-fold dilutions of serum samples or mAbs in duplicate in a total volume of 150 μl for 1 hr at 37°C in 96-well flat-bottom poly-L-lysine-coated culture plates (Corning Biocoat). Cells were suspended using TrypLE express enzyme solution (Thermo Fisher Scientific) and immediately added to all wells (10,000 cells in 100 μL of growth medium per well). One set of 8 control wells received cells + virus (virus control) and another set of 8 wells received cells only (background control). After 66-72 hrs of incubation, medium was removed by gentle aspiration and 30 μL of Promega 1X lysis buffer was added to all wells. After a 10 minute incubation at room temperature, 100 μL of Bright-Glo luciferase reagent was added to all wells. After 1-2 minutes, 110 μL of the cell lysate was transferred to a black/white plate (Perkin-Elmer). Luminescence was measured using a PerkinElmer Life Sciences, Model Victor2 luminometer. Neutralization titers are the serum dilution (ID50/ID80) or mAb concentration (IC50/IC80) at which relative luminescence units (RLU) were reduced by 50% and 80% compared to virus control wells after subtraction of background RLUs. Maximum percent inhibition (MPI) is the % neutralization at the lowest serum dilution or highest mAb concentration tested. Serum samples were heat-inactivated for 30 minutes at 56°C prior to assay.

### Surface Plasmon Resonance

Antibody binding to SARS-CoV-2 spike constructs was assessed by surface plasmon resonance on a Biacore T-200 (GE-Healthcare) with HBS buffer with 3 mM EDTA and 0.05% surfactant P-20 added. For the data presented in Figure 1H and Supplementary Figure 2, spike samples that were incubated at 4 °C, 22 °C and 37 °C for a week, were held in the sample chamber at 8 °C, 22 °C and 37 °C, respectively, during the SPR run. For the SPR data presented in Extended Data Figures 4 and 7, the sample incubated at 4 °C for a week was held in the sample chamber at 8 °C, and the samples incubated at 22 °C and 37 °C for a week, were held in the sample chamber at 22 °C during the run. The flow cell was maintained at 25 °C, thus all binding assays were performed at 25 °C. Antibodies captured on a CM5 chip coated with amine-coupled human Anti-Fc (8000RU) were assayed by SARS-CoV-2 spike at 200 nM. The surface was regenerated between injections with 3 M MgCl2 solution for 10 s at 100 μl/min. Sensorgram data were analyzed using the BiaEvaluation software (GE Healthcare).

### Cryo-EM

Purified SARS-CoV-2 spike preparations were diluted to a concentration of ~1 mg/mL in 2 mM Tris pH 8.0, 200 mM NaCl and 0.02% NaN3. A 2.5-μL drop of protein was deposited on a CF-1.2/1.3 grid that had been glow discharged for 30 seconds in a PELCO easiGlow™ Glow Discharge Cleaning System. After a 30 s incubation in >95% humidity, excess protein was blotted away for 2.5 seconds before being plunge frozen into liquid ethane using a Leica EM GP2 plunge freezer (Leica Microsystems). Frozen grids were imaged in a Titan Krios (Thermo Fisher) equipped with a K3 detector (Gatan).

## Data Availability

The datasets generated the current study are available from the corresponding author on reasonable request.

---

**Extended Data Figure 1.**
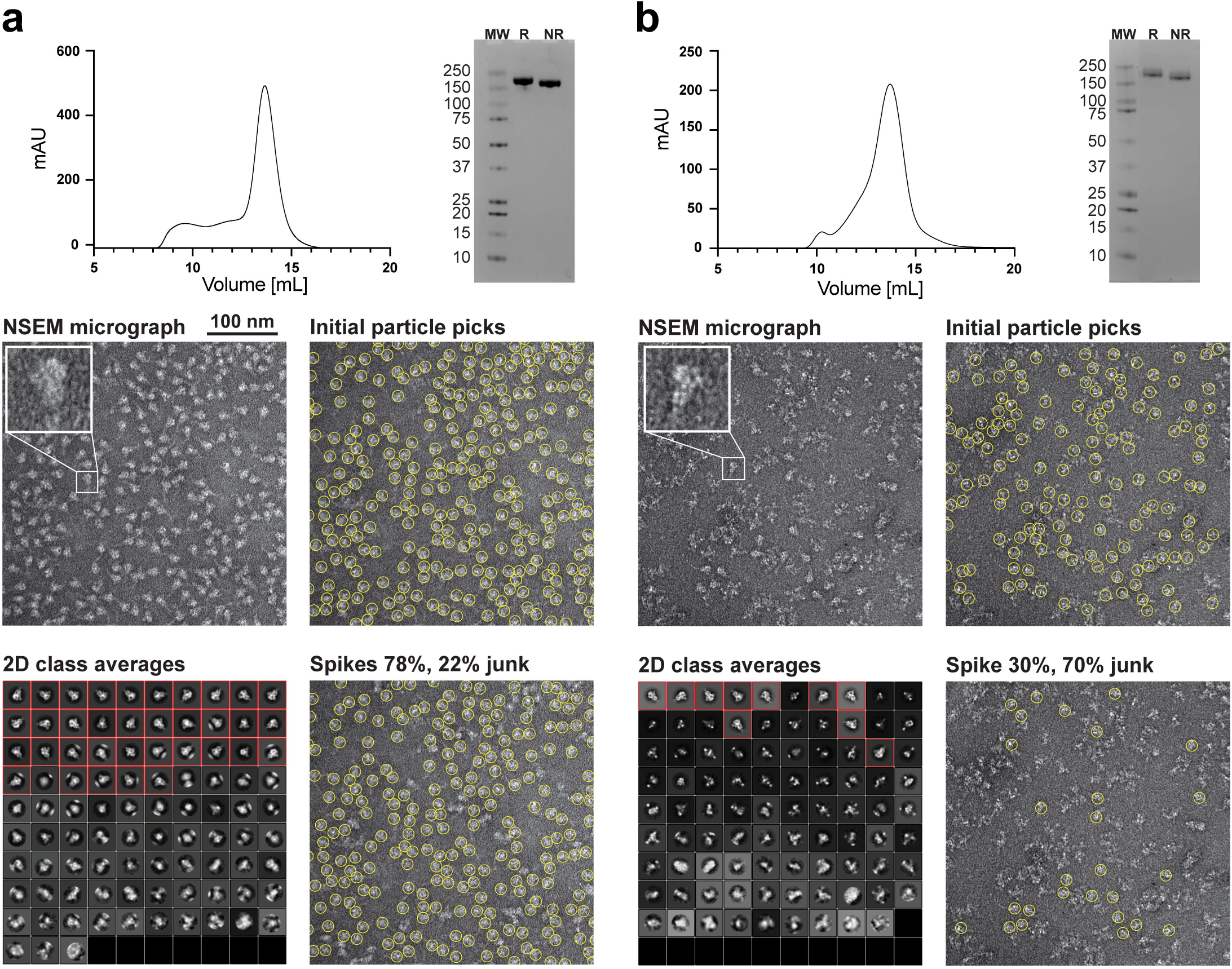
Purification and quality control of the SARS-CoV-2 S protein ectodomain. **(a and b)** Two representative spike preparations that together highlight the role of NSEM in discriminating between good **(a)** and bad **(b)** spike preparations that otherwise appear to be of similar quality by SEC (top, left) and SDS-PAGE (top, right). Representative micrographs from each prep are shown, middle left. Protein appears as white blobs on gray background. Insets show a single kite-shaped spike particle enlarged. At the middle right, automatic particle picking is shown as yellow circles superimposed on the micrograph. Sets of ~20,000 initial particles picks are subjected to automated 2D classification to group together and average particles with similar features into discrete classes, bottom left. In the 2D class averages for each sample, the classes that contain the SARS-Cov-2 S ectodomain (shown within red boxes) can be clearly distinguished from classes that contain junk. The final spike picks come from the particles contained within the indicated classes and their total number provide an estimate of the ratio between the SARS-CoV-2 S ectodomain and junk seen in the NSEM sample.

**Extended Data Figure 2.**
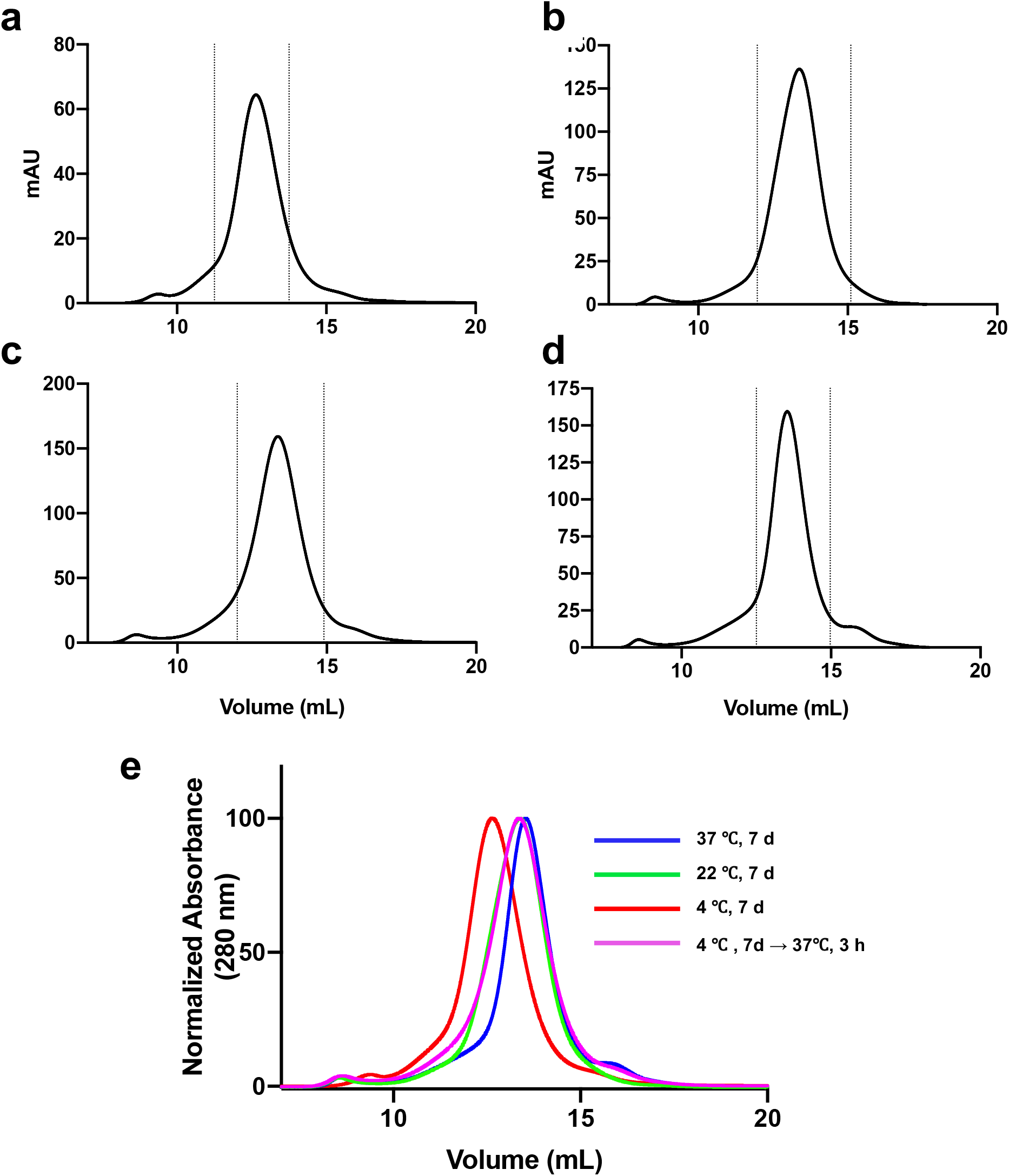
Size Exclusion Chromatography of SARS-CoV-2 S ectodomain incubated at different temperatures. SEC profiles run on a Superose 6 increase 10/300 column for SARS-CoV-2 S ectodomain samples that were incubated at **(a)** 4 °C for one week, **(b)** 22 °C for one week. **(c)** 37 °C for one week, and **(d)** 4 °C for one week and moved to 37 °C for 3 hours prior to the experiment. **(e)** Overlay of SEC plots normalized to allow better visualization of peak shifts.

**Extended Data Figure 3.**
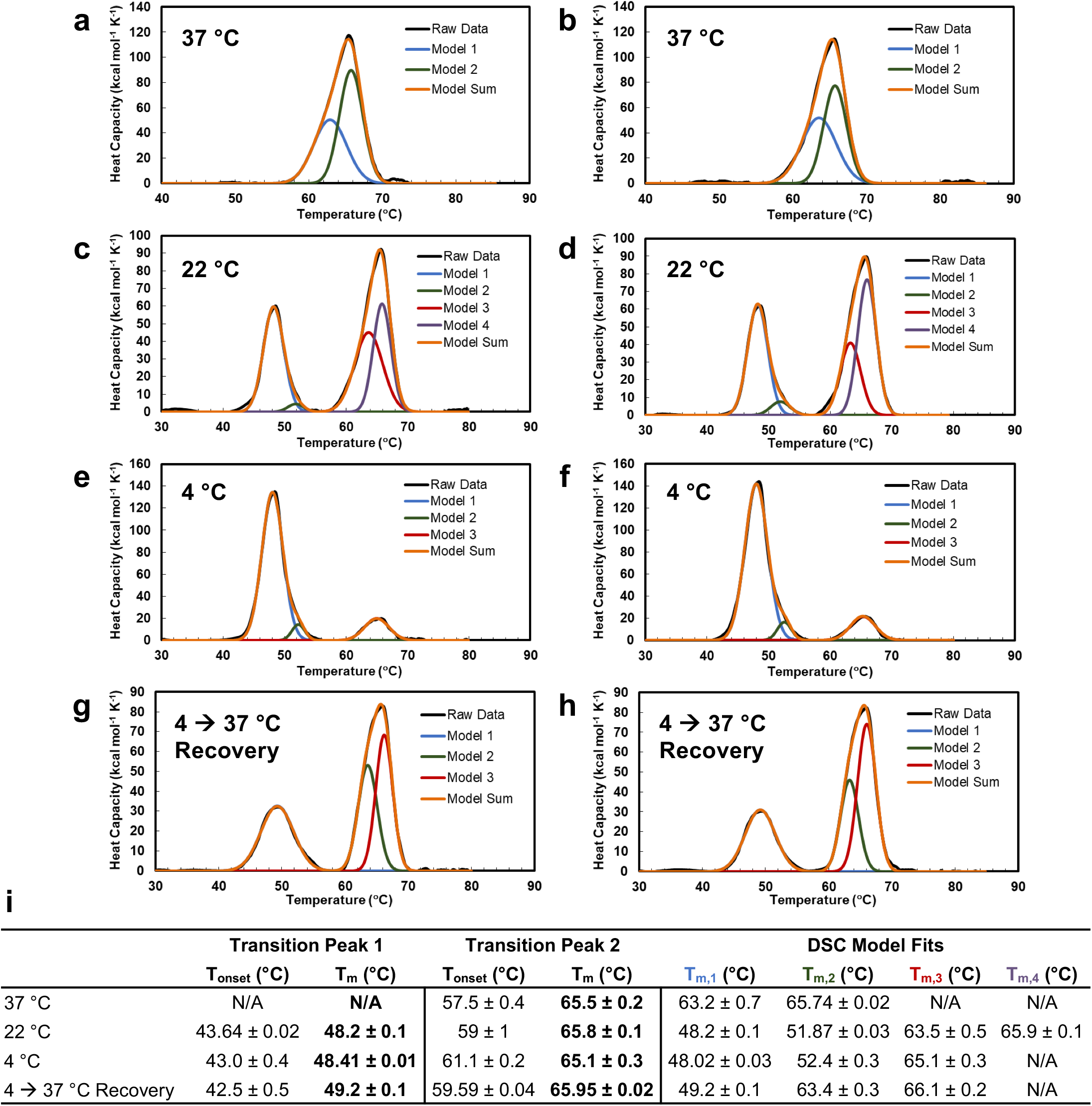
Thermostability of the SARS-CoV-2 spike ectodomain stored at different temperatures. Representative thermal denaturation profiles of the SARS-CoV-2 spike ectodomain after 1 week incubations at (**a,b**) 37 °C, (**c,d**) 22 °C, (**e,f**) 4 °C, and (**g,h**) 4 °C followed by 37 °C for 3 hours. Profiles and transition parameters (**I**) were obtained by DSC and analyzed as described in Methods. Raw data (black) was best fit with two or more Gaussian transition models (T_m,1-4_) (blue, green, red, purple). The peak observed at ~65 °C was best fit with two Gaussian transition models suggesting a complex unfolding mechanism. Data shown are the mean and range from two experimental replicates.

**Extended Data Figure 4.**
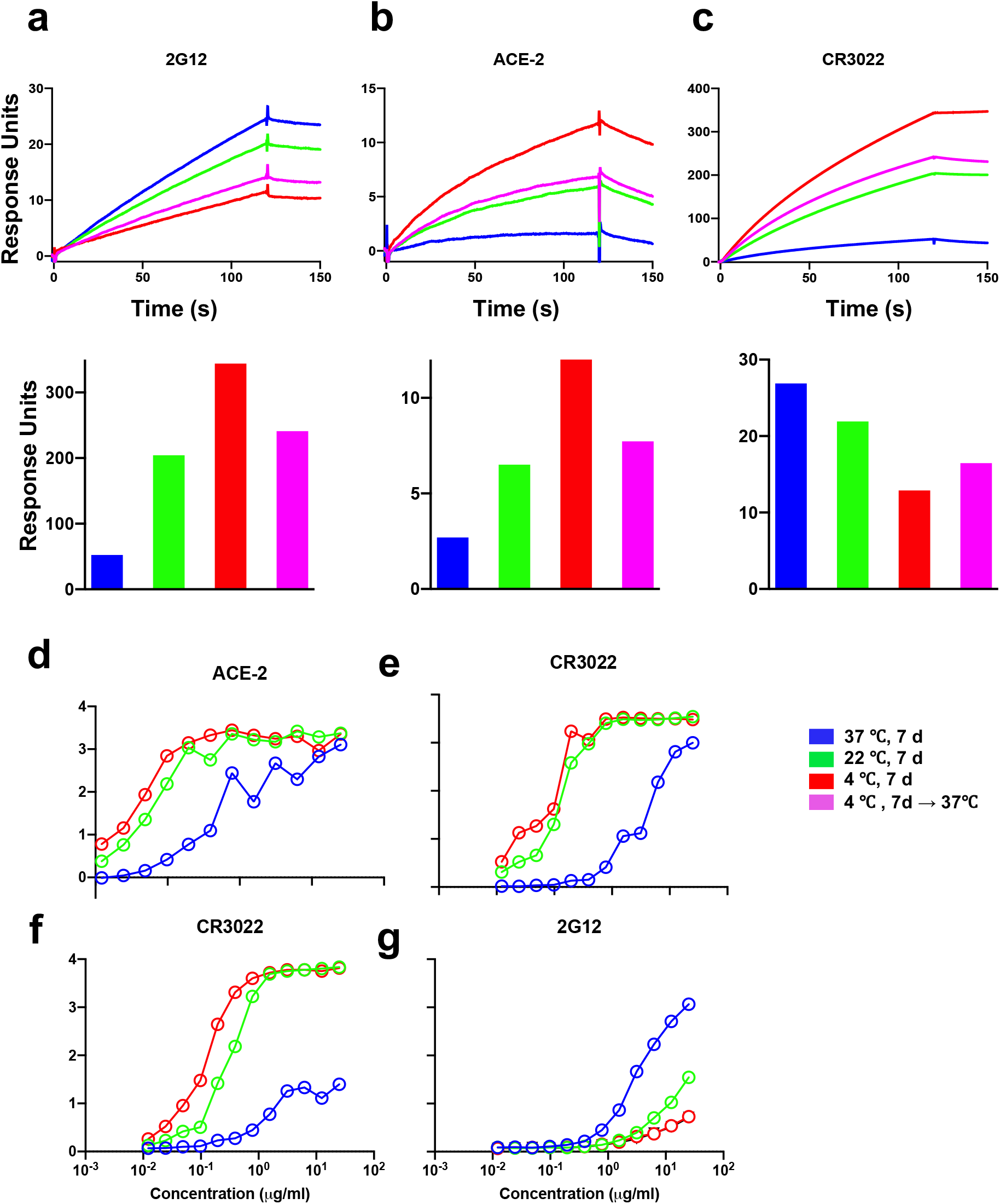
Changes in antigenicity of SARS-CoV-2 S ectodomain incubated at different temperatures. **(a-c)** SPR profiles showing binding of **(a)** ACE-2, **(b)** RBD-directed antibody, CR3022, and **(c)** S2 glycan-directed antibody 2G12 to spike samples incubated for 1 week at either 37 °C (blue), 22 °C (green) or 4 °C (red). Binding to spike sample first incubated at 4 °C for 1 week, then moved to 37 °C for 3 hours prior to the experiment is shown in magenta. **(d-g)** ELISA binding profiles showing binding of **(d)** ACE-2, **(e)** RBD-directed antibody, CR3022, in a format where antibody was coated on the plate (see methods) and **(f)** CR3022 and **(g)** 2G12, in a format where spike was captured on a strep-coated plate (see methods).

**Extended Data Figure 5.**
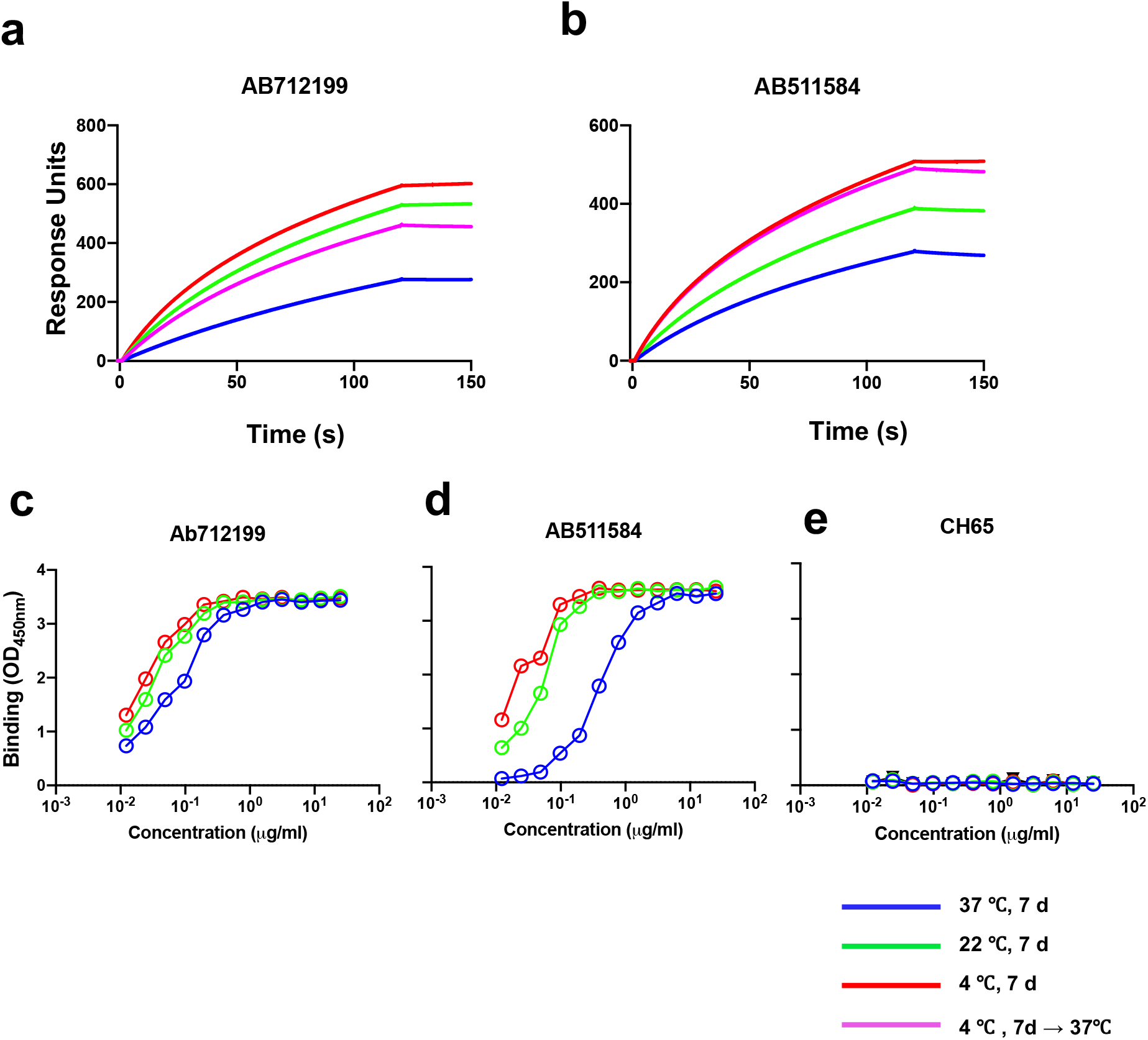
Antigenic response of SARS-CoV-2 S ectodomain incubated at different temperatures to antibodies elicited from convalescent patient sera. **(a-b)** SPR binding profiles showing binding of **(a)** RBD-directed antibody, AB712199, **(b)** S2-directed antibody, AB511584 to spike samples incubated for 1 week at either 37 °C (blue), 22 °C (green) or 4 °C (red). Binding to spike sample first incubated at 4 °C for 1 week, then moved to 37 °C for 3 hours prior to the experiment is shown in magenta. **(c-e)** ELISA binding profiles showing binding of **(c)** RBD-directed antibody, AB712199, **(d)** S2-directed antibody, AB511584, or **(e)** influenza HA-directed antibody CH65 (control) to spike samples incubated for 1 week at either 37 °C (blue), 22 °C (green) or 4 °C (red).

**Extended Data Figure 6.**
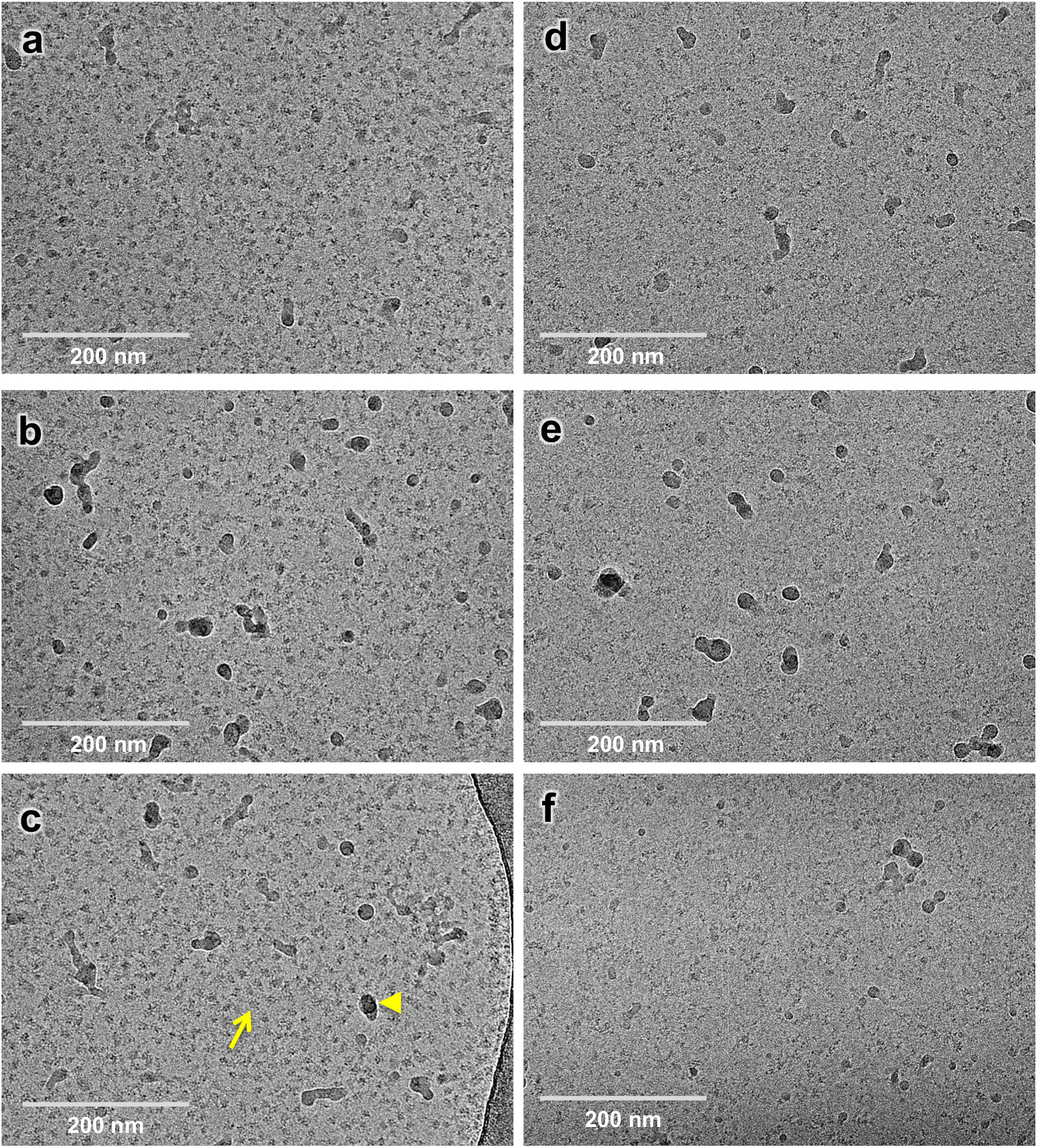
Effect of SARS-CoV-2 ectodomain spike sample storage on cryo-EM specimen preparation. **(a-c)** Representative cryo-EM micrographs of a SARS-CoV-2 S ectodomain sample that was flash frozen immediately after purification and stored in −80 °C, then thawed rapidly and incubated for ~ 5 min at 37 °C immediately prior to grid preparation. Cryo-EM images are low contrast, and the desired spike particles appear as medium gray spots (*e.g.* arrow) on a light gray background. Dark gray or black spots are slight ice contamination (*e.g* arrowhead). These panels on the left show an excellent distribution of discrete spike particles. **(d-f)** Representative cryo-EM micrographs of SARS-CoV-2 S ectodomain samples that were stored for ~1 week at 4 °C prior to grid preparation. Compared to the panels of the left, these panels on the right show a sparse field-of-view with very few intact spike particles visible. A similar spike concentration (~1 mg/ml) was used to freeze all the samples. Micrographs were collected on a Titan Krios microscope with a Gatan K3 camera.

**Extended Data Figure 7.**
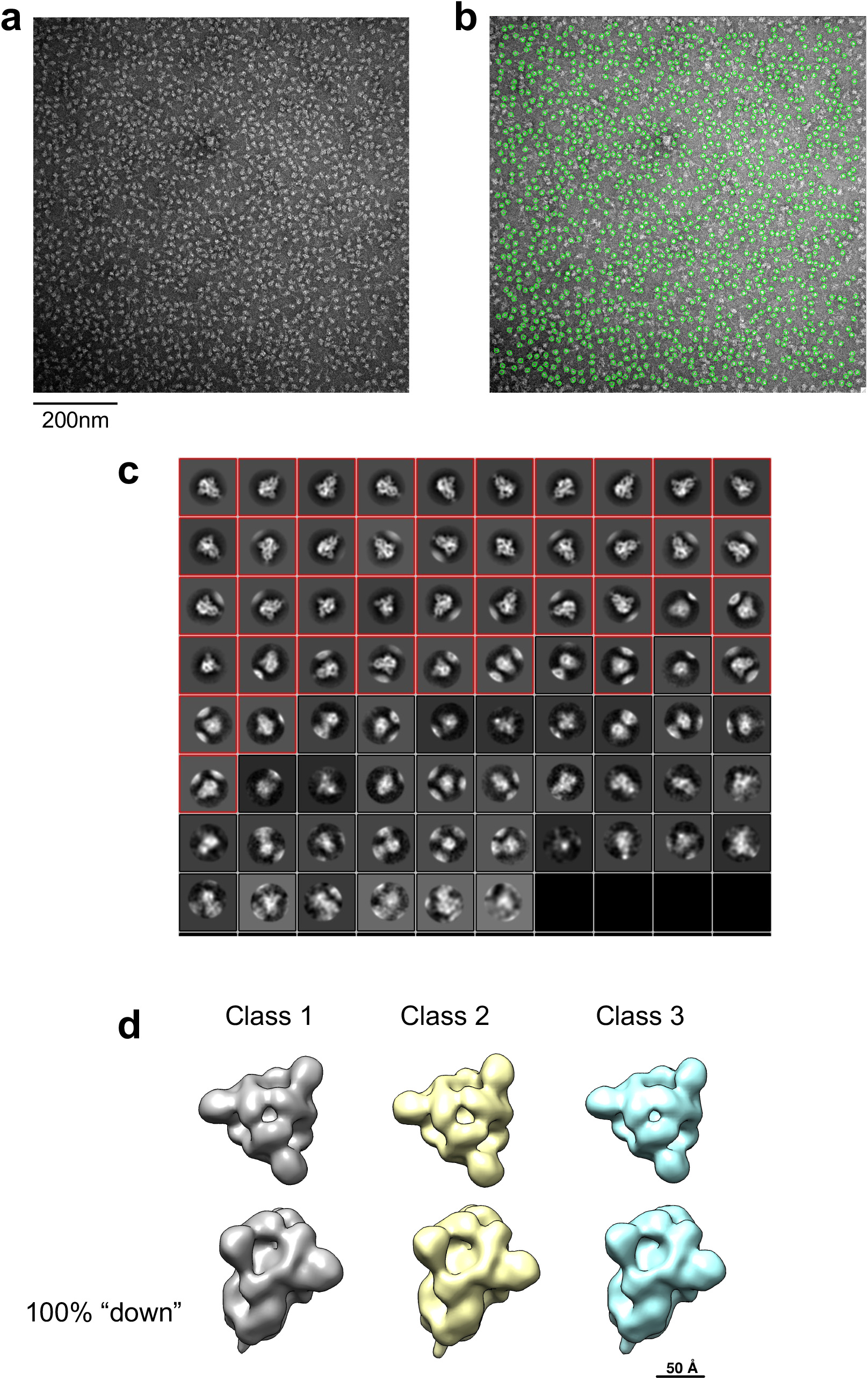
NSEM workflow for rS2d-HexaPro. **(a)** Representative NSEM micrograph **(b)** Representative NSEM micrograph showing particle picks in green **(c)** 2D class averages; the particles in the classes marked with a red box were taken forward to the next steps on the analysis **(d)** 3D classes showing top views in the top row and side views in the bottom row. 30,000 particles were used for 3D classification. These were separated into 3 3D classes that were reconstructed using C1 symmetry. Only all-RBD-down classes were observed. Also see Supplementary Movie S1 that shows the residual movement in the RBDs despite being locked down by a RBD-to-S2 disulfide.

**Supplementary Table 1.** Thermostability of the SARS-CoV-2 spike ectodomain stored at different temperatures measured by DSF. Inflection temperatures from DSF plots shown in Figure 1F, expressed as averages ± standard deviation, N=5.

| Storage Temperature | $T_1^i$ | $T_2^i$ | $T_3^i$ |
| --- | --- | --- | --- |
| 37 °C | 53.7 ± 1.1 | 65 ± 0.2 | 74.8 ± 0.2 |
| 22 °C | 49.9 ± 0.2 | 58.6 ± 0.3 | 74.2 ± 0.2 |
| 4 °C | 49.9 ± 0.4 | 57.5 ± 0.3 | 71.9 ± 0.2 |
| 4->37 °C Recovery | 52.2 ± 0.5 | 58.8 ± 0.5 | 74.1 ± 0.3 |

**Supplementary Table 2.** Thermostability of the SARS-CoV-2 spike ectodomain stored at different temperatures measured by DSC. Table of melting temperatures, T_m_, with values expressed as mean ± range, N=2.

| Storage Temp. | Transition Peak 1<br>$T_m$ (°C) | Transition Peak 2<br>$T_m$ (°C) |
| --- | --- | --- |
| 37 °C | N/A | 65.5 ± 0.2 |
| 22 °C | 48.2 ± 0.1 | 65.8 ± 0.1 |
| 4 °C | 48.41 ± 0.01 | 65.1 ± 0.3 |
| 4 → 37 °C Recovery | 49.2 ± 0.1 | 65.95 ± 0.02 |

**Supplementary Figure 1.**
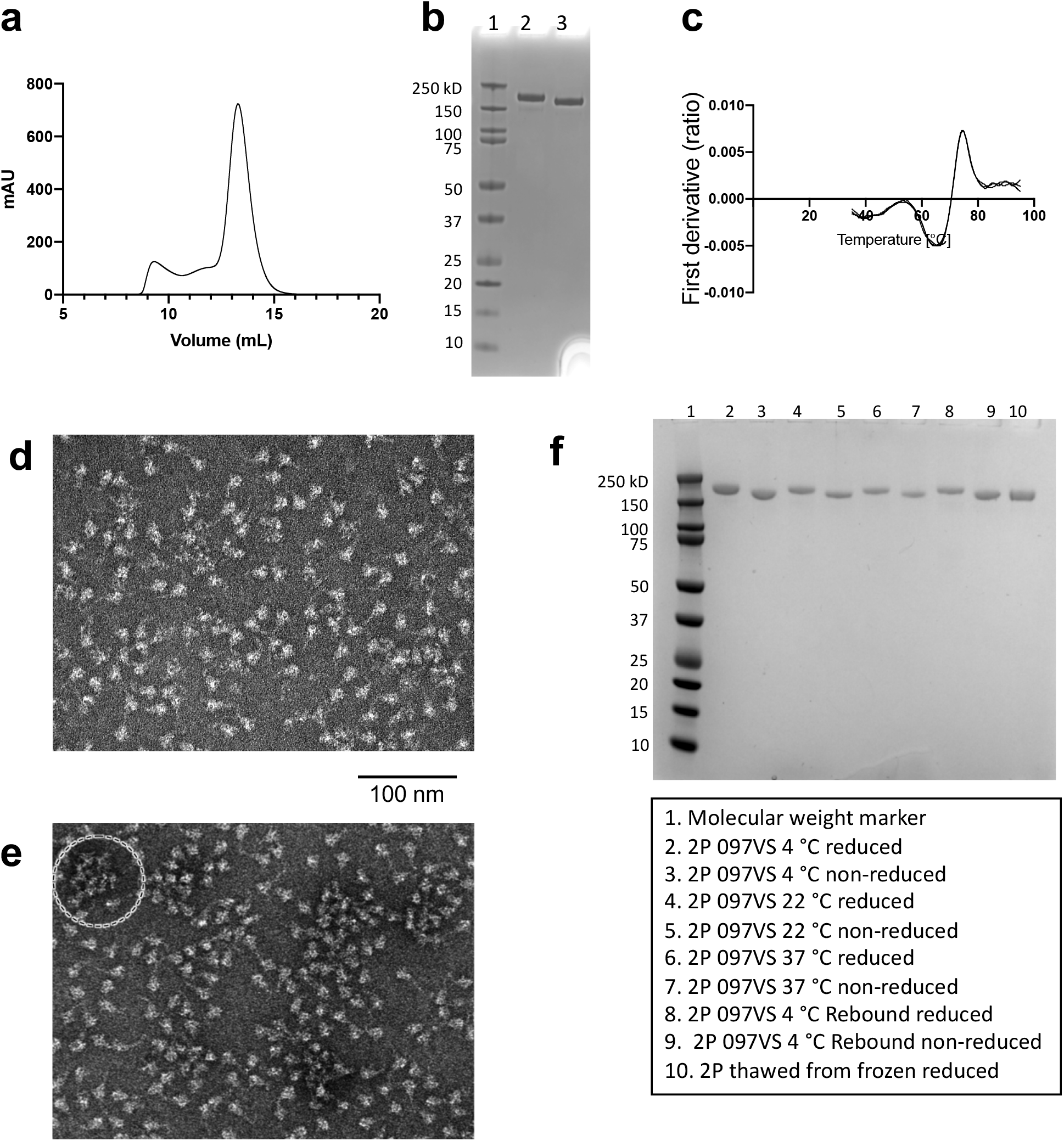
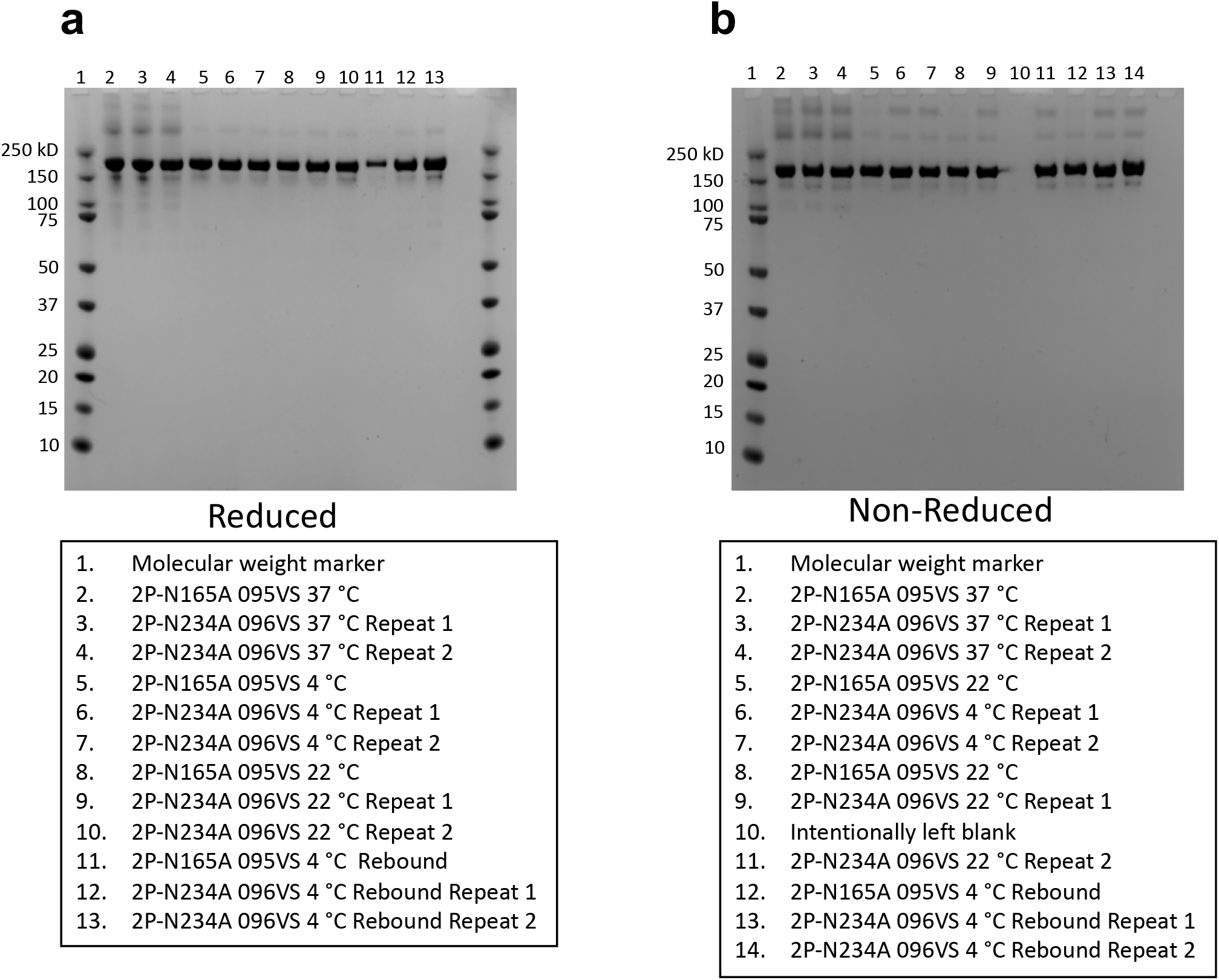
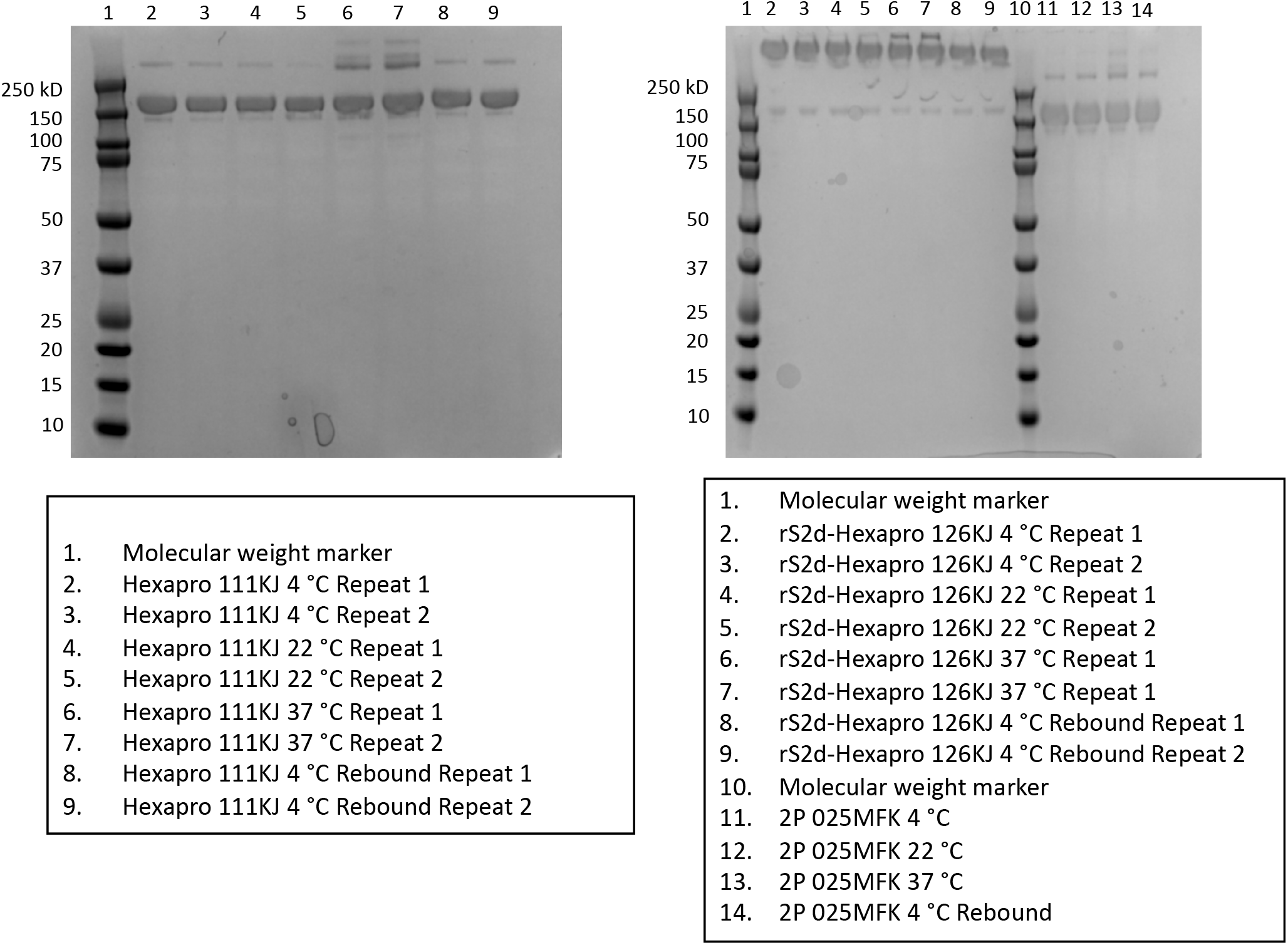
SDS-PAGE and NSEM analysis of spike proteins incubated at different temperatures. **(a)** SEC profile, **(b)** SDS-PAGE analysis (lane 1= molecular weight marker, lane 2= reducing conditions, lane 3= non-reducing conditions) and **(c)** DSF profile of a freshly purified sample of the 2P spike. **(d)** NSEM micrograph of 2P spike after storage at 22 °C for one week. **(e)** NSEM micrograph of 2P spike after storage at 4 °C for one week followed by recovery at 37 °C for 6 days. The circle indicates spike aggregation visible in the micrograph. **(f)** SDS-PAGE analysis of the 2P spike incubated at different temperatures. The legend under the gel lists the sample loaded in each lane. The sample description follows this order-construct name (e.g. 2P), lot # (e.g. 097VS), temperature at which sample was incubated for 1 week (e.g. 4 °C). Samples that were incubated first for 1 week at 4 °C, then moved to 37 °C are labeled as “Rebound”. Reduced and non-reduced sampled are noted. 3 μg sample was loaded in each well. **(a)** SDS-PAGE under reducing conditions and **(b)** SDS-PAGE under non-reducing conditions. The legend below each gel lists the sample loaded in each lane. The sample description follows this order-construct name (e.g. 2P-N165A), lot # (e.g. 095VS), temperature at which sample was incubated for 1 week (e.g. 4 °C). Samples that were incubated first for 1 week at 4 °C, then moved to 37 °C hours are labeled as “Rebound”. 3 μg sample was loaded in each well. SDS-PAGE under non-reducing conditions. The legend below each gel lists the sample loaded in each lane. The sample description follows this order-construct name (e.g. Hexapro), lot # (e.g. 111KJ), temperature at which sample was incubated for 1 week (e.g. 4 °C). Samples that were incubated first for 1 week at 4 °C, then moved to 37 °C are labeled as “Rebound”. 10 μg sample was loaded in each well.

**Supplementary Figure 2.**
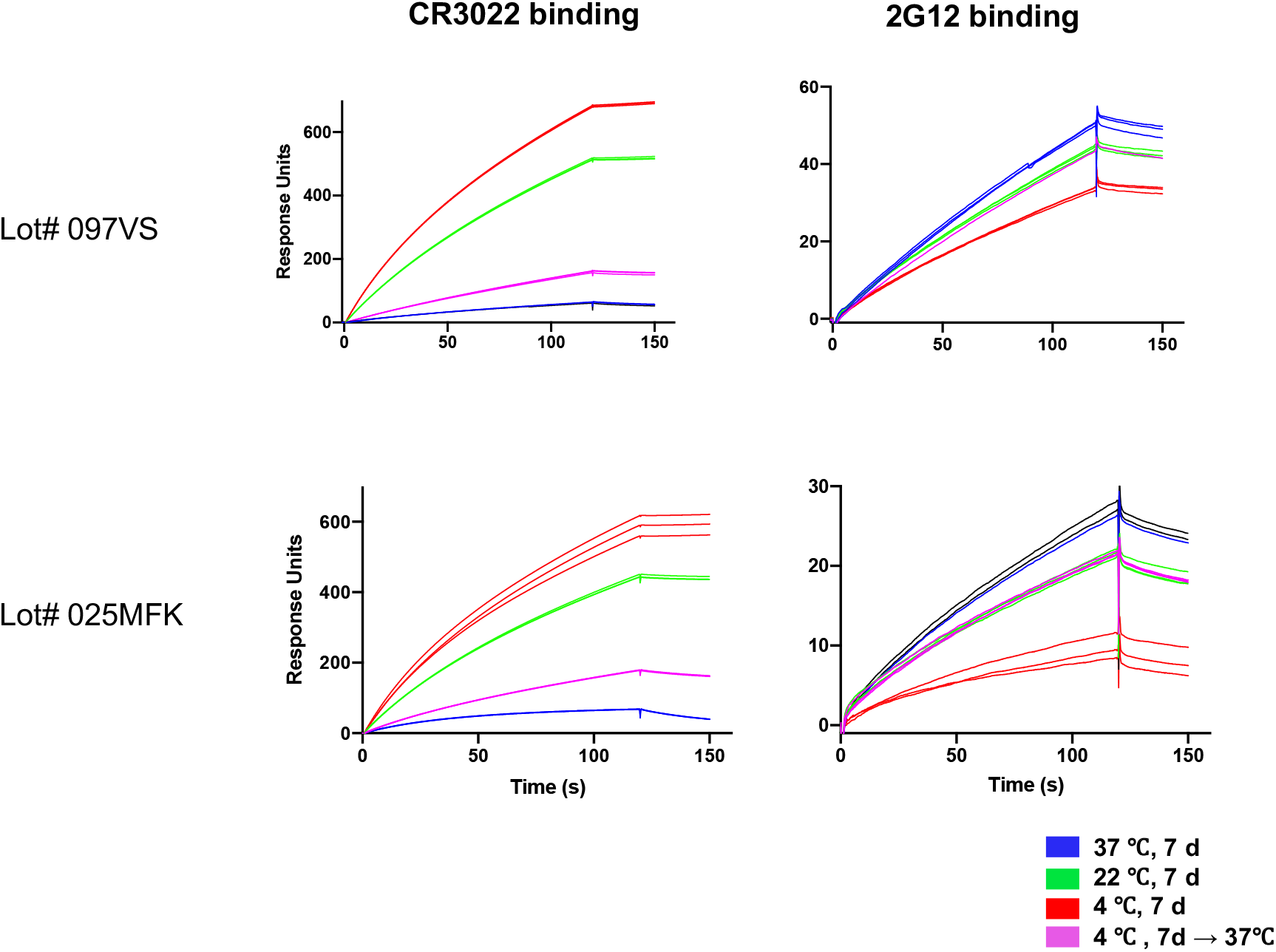
Effect of storage temperature on 2P spike antigenicity measured with two different protein lots. Antibody CR3022 IgG (left) and 2G12 IgG (right) binding to spike stored at different temperatures measured by SPR. Data for spike samples measured after a 1-week incubation at 37, 22, and 4 °C, are shown in blue, green, and red respectively; sample stored 1 week at 4 °C and then incubated for 6 hours at 37 °C shown in magenta. During the SPR run the sample chamber was maintained at temperatures of 37 °C, 22 °C and 8 °C, for the 37 °C, 22 °C and 4 °C incubated samples, respectively. The binding experiments were carried out at 25 °C.

**Supplementary Figure 3.**
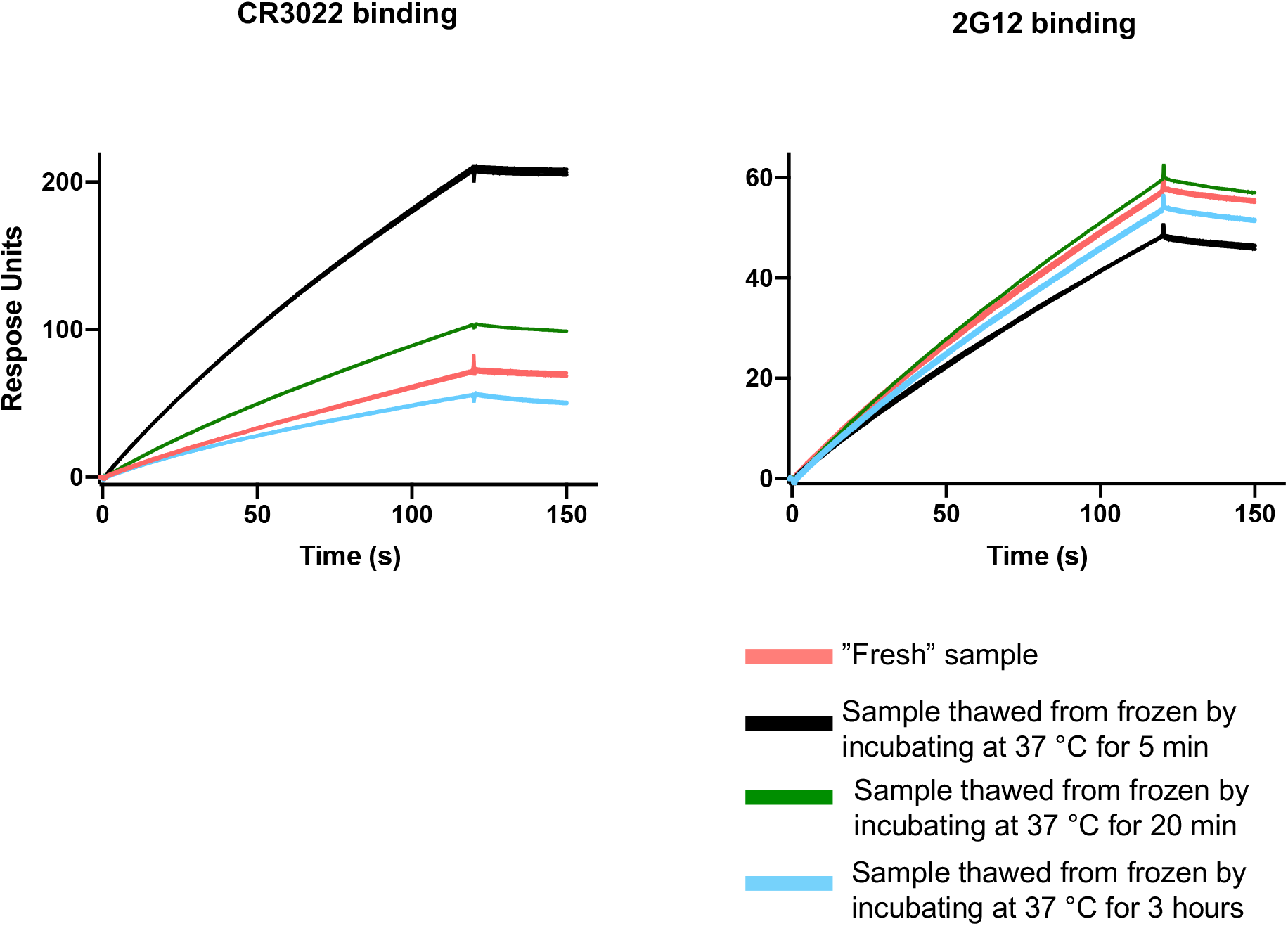
CR3022 and 2G12 binding to freshly purified spike and spike that has been flash frozen then thawed. Antibody CR3022 IgG (left) and 2G12 IgG (right) binding to spike that was either freshly purified or flash-frozen in liquid N2, then thawed by incubating at 37 °C for different periods of time.

**Supplementary Figure 4.**
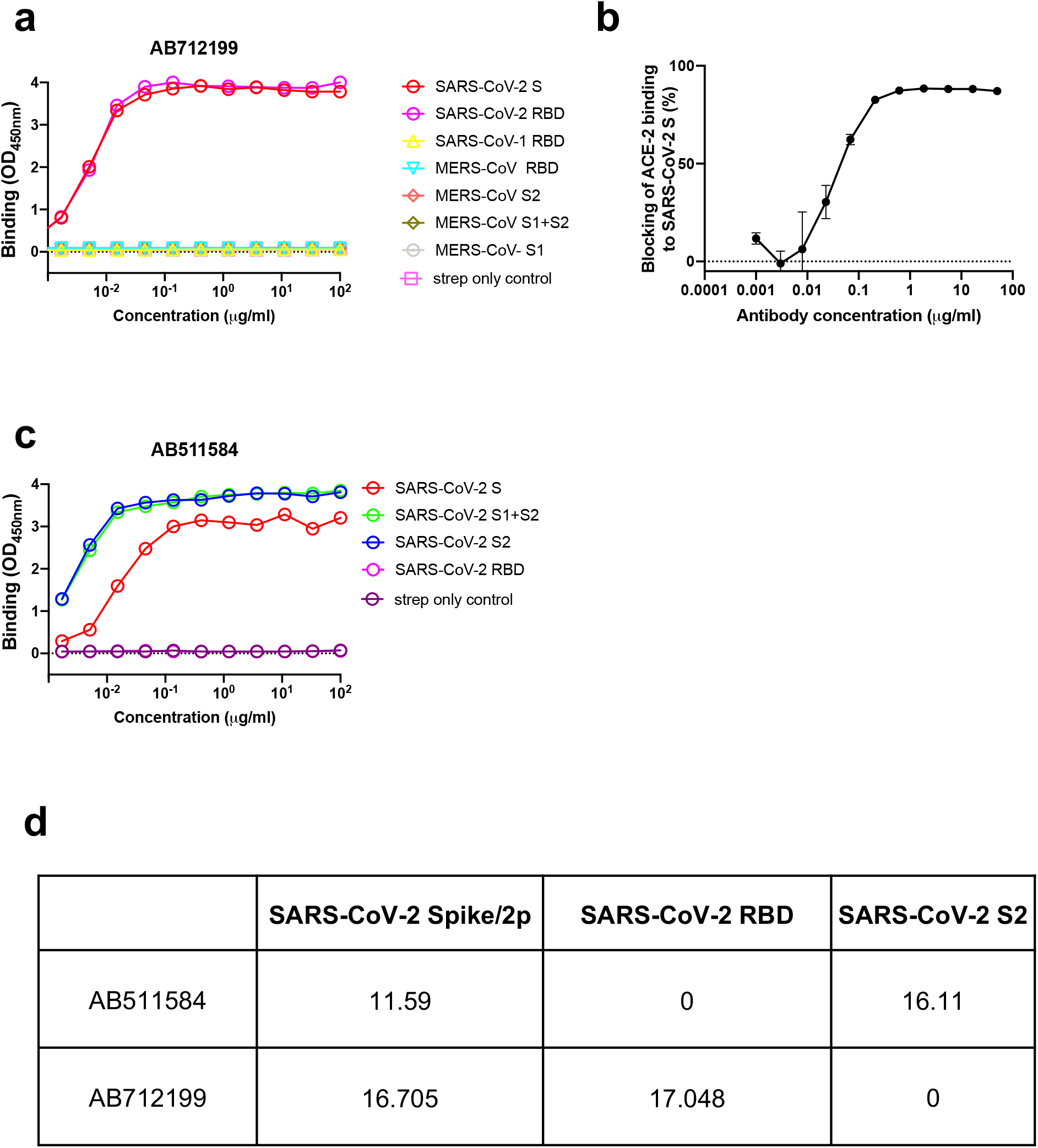
Characterization antibodies isolated from COVID-19 convalescent donors by ELISA. **(a)** Epitope mapping of antibody AB712199 shows that the antibody binds the SARS-CoV-2 RBD region, (**b**) Blocking of ACE-2 binding to SARS-CoV-2 S ectodomain by AB712199, **(c)** Epitope mapping of antibody AB511584 shows that the antibody binds the SarS-CoV-2 S2 region **(d)** Summary of epitope mapping from ELISA showing log of the area under the curves.

**Supplementary Figure 5.**
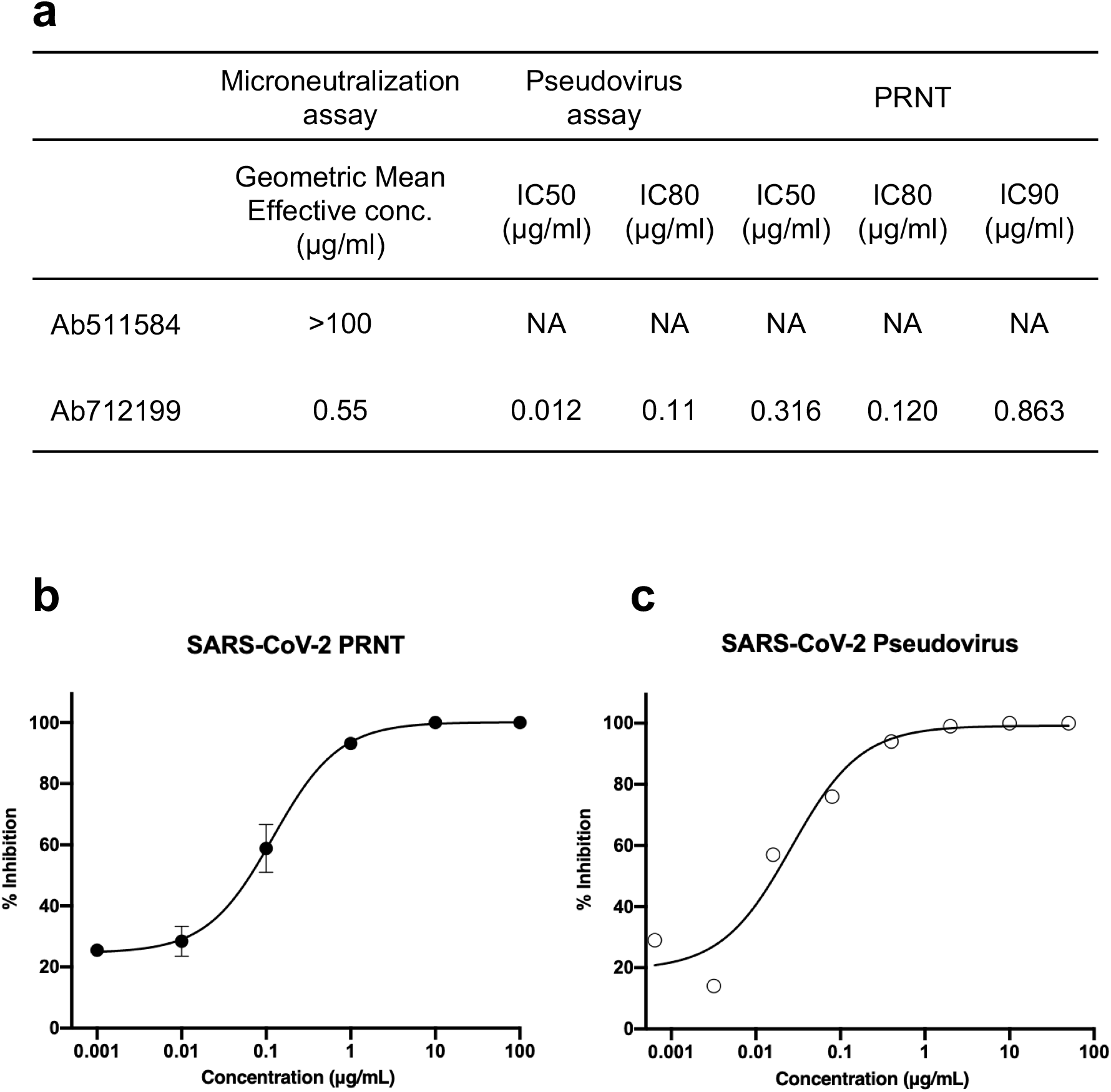
Characterization antibodies isolated from COVID-19 convalescent donors in neutralization assays. **(a)** Summary of neutralization profiles of antibodies AB511584 and AB712199 by three different assays. (**b**) Neutralization of SASR-CoV-2 by AB712199 measured using plaque reduction neutralization test. **(c)** Neutralization of SASR-CoV-2 by AB712199 measured using a pseudovirus assay.

## Notes

### Competing Interest Statement

The authors have declared no competing interest.

### Summary of Updates

1. Figures have been re-organized. 2. New data showing characterization of two antibodies isolated from convalescent COVID-19 patients added. 3. Methods updated.

